# t2pmhc: A Structure-Informed Graph Neural Network to Predict TCR-pMHC Binding

**DOI:** 10.64898/2026.02.27.708137

**Authors:** Mark Polster, Josua Stadelmaier, Elias Ball, Jonas Scheid, Jens Bauer, Annika Nelde, Manfred Claassen, Marissa Dubbelaar, Juliane S. Walz, Sven Nahnsen

## Abstract

Mapping of T cell receptors (TCRs) to their cognate MHC-presented peptides (pMHC) is central for the development of precision immunotherapies and vaccine design. However, accurate prediction of TCR affinity to peptide antigens remains an open challenge. Most approaches rely solely on sequence information, although increasing evidence suggests that TCR-pMHC binding is primarily determined by three-dimensional structural interactions within the entire TCR-pMHC complex. Consequently, sequence-based methods often fail to generalize to peptides not included in the training data (unseen peptides).

Here we introduce t2pmhc, a structure-based graph neural network framework for predicting TCR-pMHC binding using predicted structures of the entire TCR-pMHC complex. We evaluated a Graph Convolutional Network (GCN) and a Graph Attention Network, both demonstrating improved generalization to unseen peptides compared to state-of-the-art models across a variety of public datasets. Evaluation with crystallographic structures yields high-confidence predictions, indicating that current limitations of structure-based models are largely driven by the accuracy of structure prediction. Analysis of node attention patterns in t2pmhc-GCN reveals biologically consistent patterns, assigning high attention to the peptide and the CDR3 regions. Within the peptide sequence, canonical MHC anchor residues are consistently downweighted, whereas potential TCR-binding residues are upweighted.

These findings establish t2pmhc as a structure-informed framework for robust TCR-pMHC binding prediction, enabling improved generalization to unseen antigens and providing a foundation for integrating TCR repertoire sequencing into vaccine design and immunotherapy.

## Introduction

Since the initial description of the genes encoding the T cell receptor (TCR) in 1988^1^, deciphering TCR-mediated antigen recognition has been a central objective, not only advancing basic immunology but also providing a critical foundation for the development of innovative immunotherapeutic strategies. However, reliable prediction of TCR affinity to peptide antigens remains an ongoing challenge^2,3^. T cells, key players in adaptive immunity, express TCRs, to recognize short peptide antigens presented by the Major Histocompatibility Complex (MHC) on the surface of cells. Most TCRs are composed of an α-chain and a β-chain and constitute a recognition system based on high sequence diversity that has majorly evolved to counter the hypervariability of peptide-MHC (pMHC) ligands derived from viral, bacterial or malignant antigens^4,5^. TCR diversity is generated through VJ and V(D)J recombination of the α-chain and β-chain, respectively^6^. Furthermore, both chains contain three highly diverse complementarity-determining regions (CDRs)^4^. Structurally, the CDR3 regions usually form direct contacts with the presented peptide, whereas CDR1 and CDR2 predominantly interact with the α1 and α2 MHC helices^7^.

Advancement of high-throughput TCR sequencing enabled large-scale *in silico* characterization of the TCR repertoire in both patients and healthy individuals^8–11^. When combined with established mass spectrometry-based immunopeptidome profiling of pMHCs, reliable mapping of TCRs to their cognate peptides could substantially improve target discovery by antigen prioritization and thereby broaden the scope and precision of T cell-based immunotherapeutic applications. In addition, a probabilistic assessment of TCR binding may enable monitoring T cell-mediated immune responses following immunotherapy.

TCR-pMHC binding prediction remains challenging due to i) the extreme combinatorial diversity of TCRs and pMHCs, ii) the incomplete understanding of the relevant biophysical determinants, and iii) the sparseness of the training data. Although numerous sequence-based computational approaches have been proposed^12–23^, they mostly rely solely on TCR and peptide sequence information, sometimes even restricting input to the CDR3 region and peptide alone^2^. These sequence-based models can perform well when evaluated on peptides represented in the training set (seen peptides), however, they typically fail to generalize to the more clinically relevant unseen peptide setting^3^, in which test peptides are absent from training data.

Docking of TCRs onto the pMHC is not always constrained to the canonical orthogonal orientation: substantial rotation along the peptide length as well as reversed docking geometries have been observed^4^. These findings underscore that TCR-pMHC binding is fundamentally a structural problem. Recently, structure-based approaches for TCR–pMHC binding prediction have been proposed^24–28^, many of which focusing specifically on the interaction interface^24–26^. However, recent evidence suggests that the entire TCR-pMHC complex is critical for binding, and that incorporating this full structural context can improve predictive performance^2^, which in turn motivates the incorporation of three-dimensional structural information of the entire TCR-pMHC complex in predictive models.

Despite its importance, structural data remains scarce and crystallographic samples of solved TCR-pMHC complexes are very rare, limiting the feasibility of training deep learning modes directly on crystallographic complexes.

Protein-structure predictors such as AlphaFold^29^ provide a pathway toward such structure-informed binding prediction models by enabling the *in silico* generation of three-dimensional structures of TCR-pMHC complexes. Yet, general-purpose protein structure predictors struggle with TCR-pMHC complexes as the diversity of CDR loops and presented peptides is difficult to resolve^30^. Consequently, several specialized predictors have recently been developed^31–34^. However, predicted complexes remain less reliable than crystallographic structures and confidence estimates, such as Predicted Aligned Error (PAE) and predicted Local Distance Difference Test (pLDDT), which estimate the likelihood of the correctness of the structure^35^ must be considered when utilizing predicted structures.

Here, we introduce t2pmhc, a new structure-based deep learning framework for TCR-pMHC binding prediction. t2pmhc systematically integrates the full geometry of the TCR-pMHC complex into a residue-level interaction graph encoding distance-based structural associations in two variants: a Graph Convolutional Network (t2pmhc-GCN) and a Graph Attention Network (t2pmhc-GAT). t2pmhc employs attention to capture key binding determinants and gives insights into residue-level attention patterns. In a benchmark against state-of-the-art methods across multiple independent test datasets, t2pmhc shows improved generalization in the challenging unseen peptide setting while maintaining competitive performance on seen peptides. By integrating full-complex structural information into graph-based learning, t2pmhc provides a framework for incorporating TCRs into peptide prioritization, immunomonitoring, and other immunotherapeutic applications, particularly in settings involving unseen peptides.

## Methods

### Sequencing Data

TCR-pMHC binders were retrieved from publicly available data repositories. Human samples with TCRα and TCRβ chains were downloaded from VDJdb^36^ (accessed 23.08.2024). The McPAS database^37^ was queried for human entries (accessed 24.09.2024) and samples were downloaded from the IEDB^38^ (accessed 24.09.2024). One sample was acquired from an own previous study^39^. To ensure non-redundancy, duplicated TCR-pMHC pairs were removed based on V/J gene usage and CDR3α/β amino acid sequence similarity.

Furthermore, MHC class II data was removed due to the small number of such samples. Samples for which no structure could be predicted were removed from the final dataset. Only pMHCs associated with more than ten binding TCRs were retained for training and downstream analysis.

### Intra-dataset-based negative dataset creation

For each peptide in the dataset, TCRs known to bind to other peptides were selected as negatives. To avoid similarities with the peptide of interest, only TCR-peptide pairs where the peptide had a Levenshtein distance greater than 3 from the query peptide were considered. For each positive observation, five negative samples were generated. Two negatives of peptides of different lengths and three negatives from peptides of similar length were sampled to mitigate TCR length bias^40^. For two peptides in the training dataset, insufficient negatives were available. In these cases, all TCRs binding to other peptides were used as negatives. The final dataset contained 20,809 positive and 82,303 negative TCR-pMHC pairs, covering 77 unique peptides and 57 MHC alleles.

### Structure Prediction

Structures of all TCR-pMHC complexes were predicted using TCRdock (v2.0.0)^33^. To efficiently predict TCR-pMHC structures, TCRdock was added to the nf-core/proteinfold^41^ pipeline in a feature branch. Structures were predicted in batches of 500 and *new_docking* was enabled to speed up the prediction. To capture structural uncertainty in TCR-pMHC complexes, the PAE was obtained from AlphaFold’s ‘model_2_ptm’ parameter via TCRdock. A TCR-pMHC-specific PAE was calculated via TCRdock as described by Bradley^33^.

### t2pmhc Architecture

The t2pmhc framework comprises structural preprocessing of predicted complexes, graph-based representation learning, and supervised classification for the t2pmhc-GCN and t2pmhc-GAT model variants.

#### Contact Map Generation

TCR-pMHC complex structures predicted using TCRdock were translated into residue-level contact maps. For each pair of amino acid residues, the Cα-Cα distance was calculated. Residues within 10 Ångström (Å) were considered in contact. The resulting binary contact maps were stored as adjacency matrices and used as the structural basis for graph construction in both t2pmhc models. This representation captures the three-dimensional organization of the TCR-pMHC complex in a format suitable for graph neural networks.

#### Graph Construction

TCR-pMHC complexes were represented as residue-level graphs G = (V, E) based on the aforementioned contact maps. Nodes v ∈ V corresponded to amino acids and edges e ∈ E captured spatial contacts derived from the contact map. Node features included amino acid type using the tcrBLOSUM^42^, hydrophobicity, formal charge, Atchley factors^43^, domain affiliation (TCRα/β, peptide, MHC, CDR3α/β, or peptide residues), and the Predicted Aligned Error (PAE) averaged over all TCR-pMHC residue pairs. Each edge was annotated with its Cα-Cα distance (in Å). For t2pmhc-GAT, edges were additionally annotated with the PAE between each residue pair. Graph construction was parallelized to process thousands of structures efficiently. t2pmhc supports saving/loading preprocessed graphs to ensure reproducibility and avoid redundant computation.

#### Model architecture

The GCN^44^ architecture comprised three stacked graph convolutional layers, each followed by batch normalization, rectified linear unit (ReLU)^45^ activation, and dropout. The GAT^46^ architecture consisted of three stacked graph attention layers. Each layer was followed by batch normalization, exponential linear unit (ELU)^47^ activation, and dropout. For both variants, after the last layer, attention-based global pooling^48^ aggregated node representations before a fully connected layer was used to compute binding probabilities. Models were trained using cross-entropy loss and optimized with Adam^49^. The t2pmhc models were written in Python^50^ (v3.11) and built using the PyTorch^51^ (v2.6.0) and PyTorch Geometric (v2.6.1) libraries for graph representation and neural network implementation.

#### Experimental setup

Twenty percent of the final dataset were randomly held out as an independent test set using a stratified split that preserved the per-peptide ratio of positive to negative samples. The remaining 80% were used for model training and hyperparameter optimization. Hyperparameters were tuned via Bayesian optimization with 5-fold cross-validation.

### Comparative Evaluation with State-of-the-Art Approaches

For benchmarking, the constructed public test set was used. In addition, the IMMREP23-solved dataset^3^ and the recently published viral dataset from the ePytope-TCR benchmarking study^52^ were included. A set of 139 human αβTCR-pMHC complexes was retrieved from the Structural T cell Receptor Database (STCRDab)^53^ (accessed: 14.07.2025). The PDB files of these crystal structures were used as input for the t2pmhc models. Resolution was used as the structure quality feature. Sequence data from these samples was retrieved and used as input for the sequence-based models. To avoid data leakage, any entries overlapping with the training data of the evaluated models were removed from each dataset before benchmarking. All datasets were further partitioned into seen and unseen based on the presence of peptides in the training data of all models (Table 1). To ensure reproducibility and facilitate automated execution of the benchmark, we implemented a Nextflow^54^ pipeline based on the nf-core^55^ template, integrating all benchmarked models and the preprocessing step.

**Table 1:** Benchmark Dataset Overview.

| Dataset | Seen |  |  | Unseen |  |  |
| --- | --- | --- | --- | --- | --- | --- |
|  | Peptides | Positive Samples | Negative Samples | Peptides | Positive Samples | Negative Samples |
| Public Test Set | 12 | 281 | 5959 |  |  |  |
| Immrep23 | 9 | 214 | 1455 | 5 | 126 | 642 |
| Epytope-TCR viral | 3 | 32 | 1882 | 4 | 177 | 2375 |
| STCRDab | 10 | 25 | 0 | 78 | 114 | 0 |

The following TCR-pMHC binding predictors were used in this benchmark:

● ERGO-II^14^ was executed via its Python implementation with minor modifications, including automated device selection and updated paths to pre-trained models. The vdjdb-based model of ERGO-II was used.
● TABR-BERT^13^ was executed using the official Docker container with pre-trained embeddings provided by the authors.
● MixTCRpred^15^ was executed using its Python implementation. Unlike the other models, it reports a percent rank score, with lower values corresponding to higher binding probability. To maintain consistency across predictors, these scores were normalized to the range [0, 1] and inverted (1 − normalized score). Since MixTCRpred employs a separate model for each peptide, predictions were performed individually for each peptide.
● MixTCRpred-pan^15^ was not publicly available at the time of writing. It was retrained on the original “pan_training_set” as described by the authors. Percent rank score was normalized and inverted as described above.

### Probability score calibration for STCRDab evaluation

The STCRDab holds crystal structures of known TCR-pMHC binders. Thus, no negatives could be retrieved from the database. As imposing negatives directly on the PDB files could have altered the docking conformation and overall geometry, no negatives were generated for the dataset. Instead, the binding probabilities were assessed directly. To make raw binding probabilities comparable between models, the binding scores were calibrated using the independent Immrep23-solved test dataset. Isotonic regression calibration^56^ was applied to fit a non-parametric, monotonically increasing mapping between raw model outputs and true outcome frequencies to address class imbalance in the dataset. Calibration models were fitted separately for each predictor and the resulting calibrators were subsequently applied to the prediction probabilities on the STCRDab test-set.

### Attention weight calculation

As GCNs do not employ the attention mechanism, there is no explicit modeling of node importances during message passing. To obtain interpretable node importances, we consider the attention-based global pooling of the final node embeddings of t2pmhc-GCN. Importantly, this node-level attention does not influence message passing within the GCN layers. To make attention weights more comparable between graphs of different sizes, attention weights were rescaled to [0,1] using Min-Max normalization.

For t2pmhc-GAT, node attention scores were rescaled to [0,1] within each structure and further normalized by in-degree to correct for connectivity bias and by domain size to enable fair comparison of attention allocation across domains of different sizes. Edge attention coefficients were softmax-normalized per node, aggregated by interacting domain, and normalized per graph to enable comparison of attention distribution across domains.

## Results

### t2pmhc, a novel Graph Neural Network-based framework to predict TCR-pMHC binding

t2pmhc leverages protein structure prediction to construct residue-level graphs of full human TCR-pMHC complexes. These graphs incorporate structural information directly into the binding prediction task. By employing a rich feature space and the entire geometry of the complex, t2pmhc captures the biophysical dependencies underlying TCR recognition (Fig. 1).

**Figure 1:**
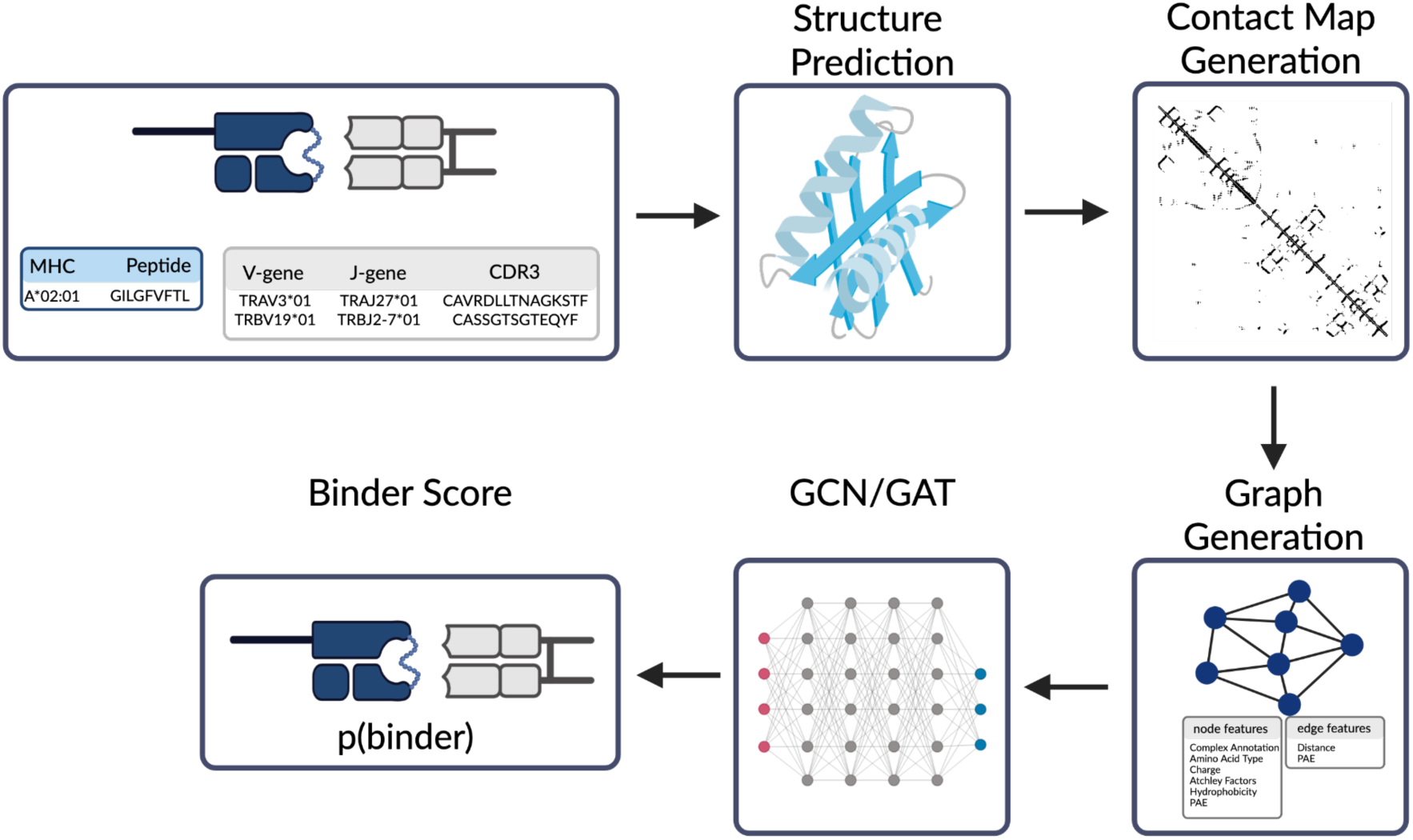
Overview of the t2pmhc framework for structure-based TCR-pMHC binding prediction. TCR α- and β-chain genes, CDR3 sequences, together with the peptide and MHC allele, are used to predict the full TCR-pMHC complex structure with TCRdock^33^. The predicted structure is translated into a Cα-Cα contact map, which is converted into a residue-level interaction graph, where nodes represent individual residues and edges encode spatial proximity within the complex. The resulting graph is processed by either a GCN or GAT. Finally, a classification head outputs a binding probability for the TCR-pMHC pair.^57^ Abbreviations: p(binder), TCR-pMHC binder probability; PAE, Predicted aligned error; MHC, major histocompatibility complex; CDR3, complementarity-determining region 3; GCN, graph convolutional network; GAT, graph attention network.

TCR-pMHC binders were aggregated from several public repositories, including VDJdb^36^, McPAS^37^ and IEDB^38^. The resulting dataset contained 77 unique peptides (Fig. 2a), and was biased toward common MHC class I alleles, with A*03:01 and A*02:01 accounting for 77.3% of all samples (Fig. 2b). The majority of peptides were derived from viral pathogens. Cytomegalovirus (CMV) and Epstein-Barr Virus (EBV) were the most prominent epitope species (Fig. 2c).

**Figure 2:**
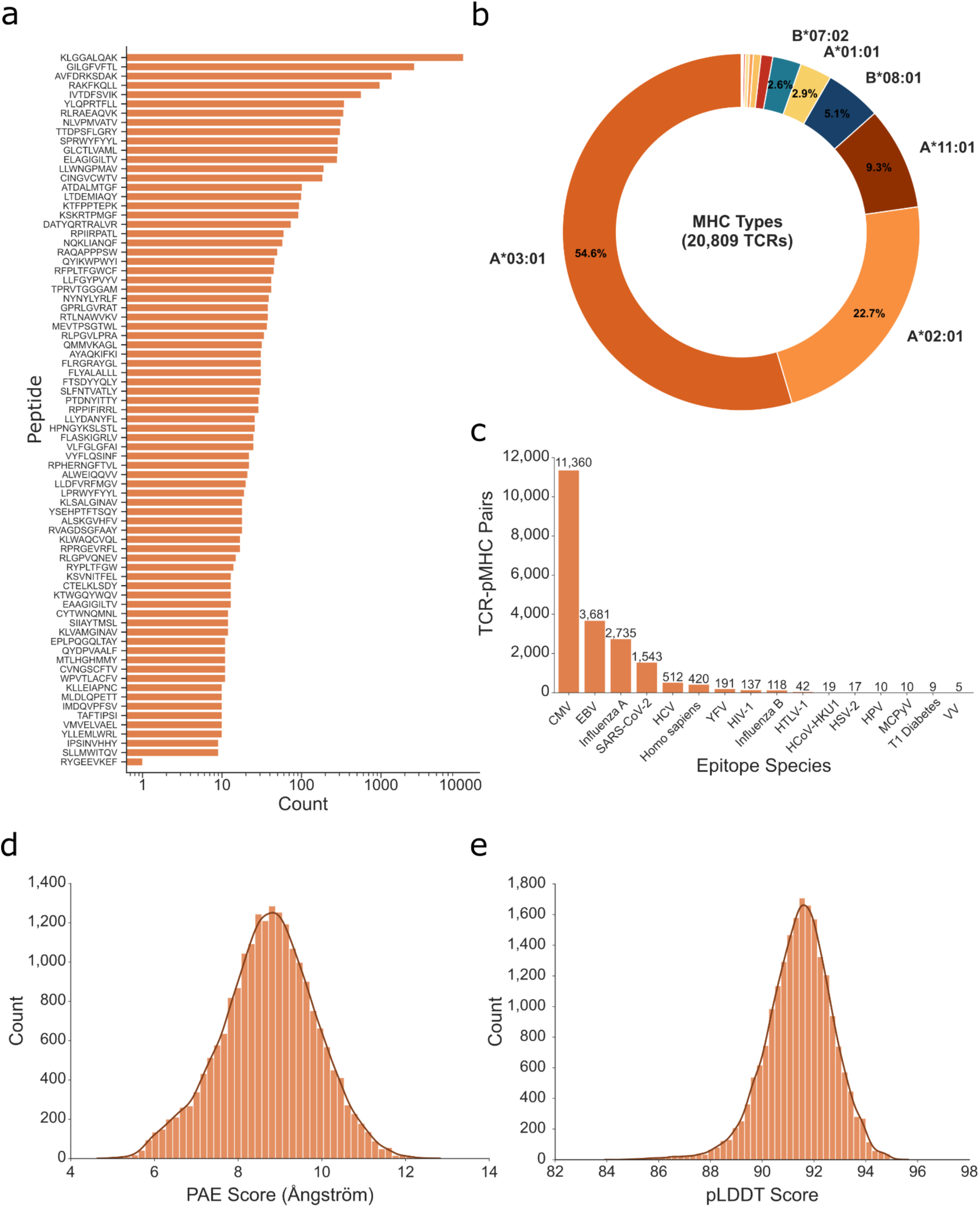
TCR-pMHC training dataset for t2pmhc. (a) Frequency of peptide sequences across the dataset. Logarithmized x-axis. (b) Distribution of MHC class I alleles across TCR-pMHC samples (20,809 TCRs). (c) Epitope species represented in the training dataset. (d) Distribution of mean predicted aligned error (PAE) across predicted TCR-pMHC complexes. (e) Distribution of local distance difference test (pLDDT) scores. Abbreviations: PAE, Predicted aligned error; pLDDT, predicted Local Distance Difference Test; CMV, Human cytomegalovirus; EBV, Epstein-Barr virus; Influenza A, Influenza A virus; SARS-CoV-2, Severe acute respiratory syndrome coronavirus 2; HCV, Hepatitis C virus; YFV, Yellow fever virus; HIV-1, Human immunodeficiency virus type 1; Influenza B, Influenza B virus; HTLV-1, Human T-lymphotropic virus type 1; HCoV-HKU1, Human coronavirus HKU1; HSV-2, Herpes simplex virus type 2; HPV, Human papillomavirus; MCPyV, Merkel cell polyomavirus; T1 Diabetes, Type 1 diabetes mellitus; VV, Varicella-zoster virus; TCR, T cell receptor; pMHC, peptide-major histocompatibility complex; MHC, major histocompatibility complex;

Structural models for all TCR-pMHC complexes were generated using TCRdock^33^. Global structural inter-residue uncertainty, assessed via the mean PAE across the entire complex, was moderate (mean PAE: 8.72 ± 1.14 Å) (Fig. 2d), reflecting the difficulty in modelling highly variable TCR-pMHC interfaces even for highly specialized tools. The predicted structures exhibited consistently high local confidence with a mean pLDDT score of 91.43 ± 1.29, indicating high residue-level confidence (Fig. 2e).

The predicted structures were subsequently translated into residue-level graphs (Fig. 3a, b).

**Figure 3:**
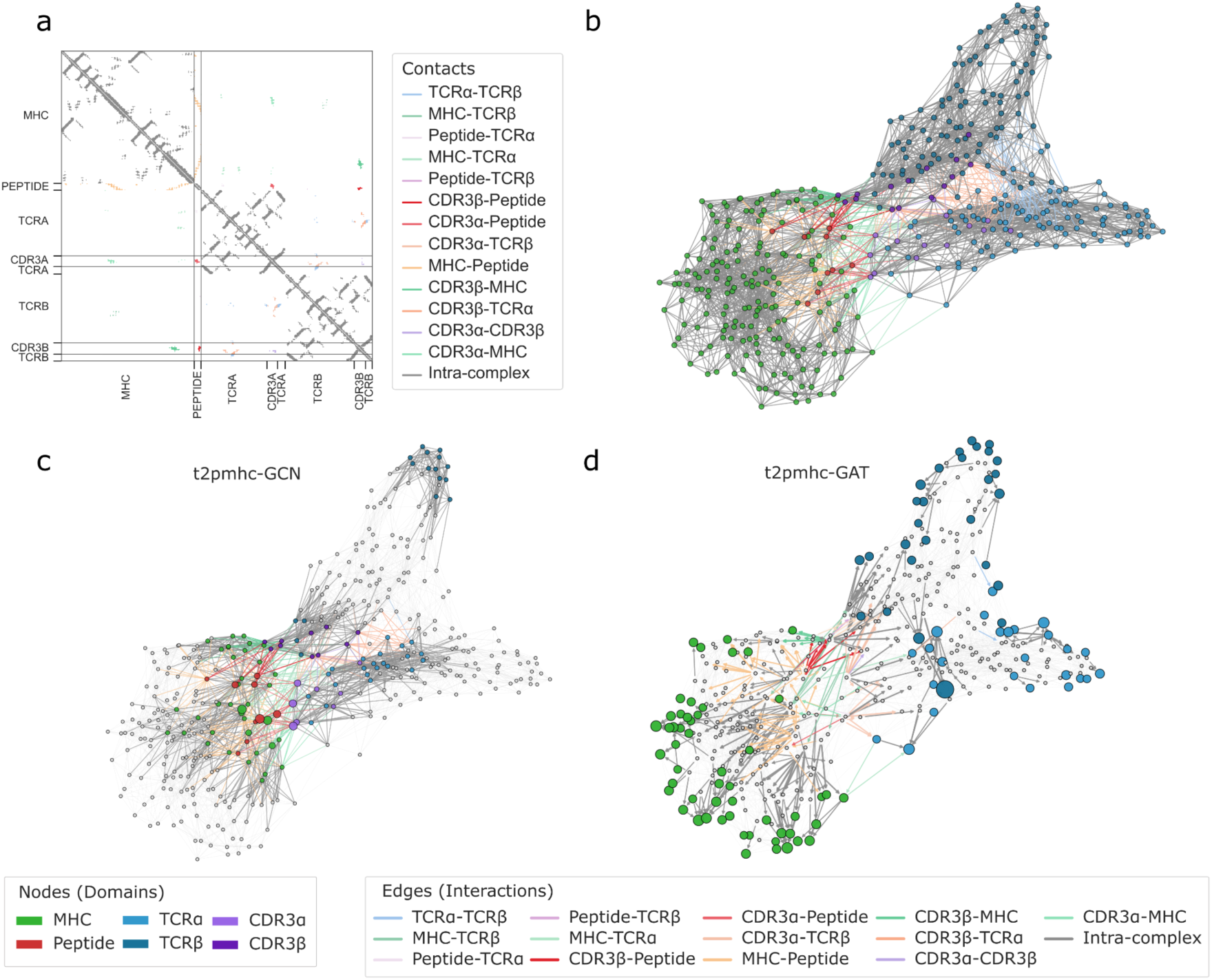
Generation of residue-level interaction graphs from predicted TCR-pMHC complex structures using a representative example of a TCR binder to the A*03:01-restricted peptide KLGGALQAK. (a) Cα-Cα contact map derived from a predicted TCR-pMHC complex, with contacts colored by interacting domains (MHC, peptide, TCRα, TCRβ, CDR3α, CDR3β). (b) Residue-level interaction graph constructed from the contact map in (a). Nodes represent individual residues and are colored by their corresponding domain. Edges represent spatial contacts between residues and are colored by the domains of the connected residues. (c) Residue-level interaction graph described in (b) after assigning node attention scores using t2pmhc-GCN. The top quartile of most important nodes is colored by its domain, while nodes with less attention are depicted in grey. Node sizes are scaled proportionally to their attention weights. (d) Residue-level interaction graph described in (b) after assigning node and edge attention scores using t2pmhc-GAT. The top 5% of most important edges are shown and self-loops are removed. The top quartile of most important nodes is colored by its domain, while nodes with less attention are depicted in grey. Node sizes are scaled proportionally to their attention weights.

### t2pmhc-GCN captures biologically relevant structural determinants of TCR-pMHC binding

t2pmhc-GCN employs attention-based global pooling, which computes attention weights over residues in the TCR-pMHC interaction graph to generate pooled graph representations^48^ (Fig. 3c). To quantify node attention, attention weights were extracted for each residue from the model for the immrep23 and public test sets. Attention weights were normalized within each TCR-pMHC complex using Min-Max normalization and the mean was computed per domain (Peptide, MHC, TCRα, TCRβ, CDR3α, CDR3β). The domain attention values were then averaged over all graphs. Across all complexes in the test sets, the peptide domain received most attention (median=0.3), followed by the CDR3 regions (CDR3α: median=0.09, CDR3β: median=0.04). In contrast, MHC (median=0.02), TCRα (median=0.002) and TCRβ (median=0.0002) domains received minimal attention (Supplementary Fig. 1a, b).

The t2pmhc-GAT model assigned most attention to the MHC, TCRα and TCRβ domains (Fig. 3d, Supplementary Fig. 1c,e).

To further investigate whether attention patterns differ between binders and negatives in t2pmhc-GCN, attention profiles per domain were stratified into quantiles by prediction probability (Fig. 4a). In both binders and negatives, the peptide domain received the highest attention across all probability quantiles, confirming that peptide features dominate model focus. Again, CDR3α and CDR3β received moderate attention, whereas the MHC, TCRα and TCRβ domains remained near zero across most quantiles. Notably, TCRβ showed a slight increase in attention in a subset of binders in the highest-confidence quantile (0.8-1.0) (Fig. 4a). When focusing on the highest-confidence quantile, a distinct redistribution of attention was observed in binders that was not present in other quantiles or in negatives. In this group, peptide attention became more variable with a lower median compared to all other quantiles, while attention assigned to TCRβ increased (Fig. 4a). CDR3β also showed a redistribution, with lower median values. Negatives did not display this pattern in any quantile; attention patterns remained comparable across all quantiles.

**Figure 4:**
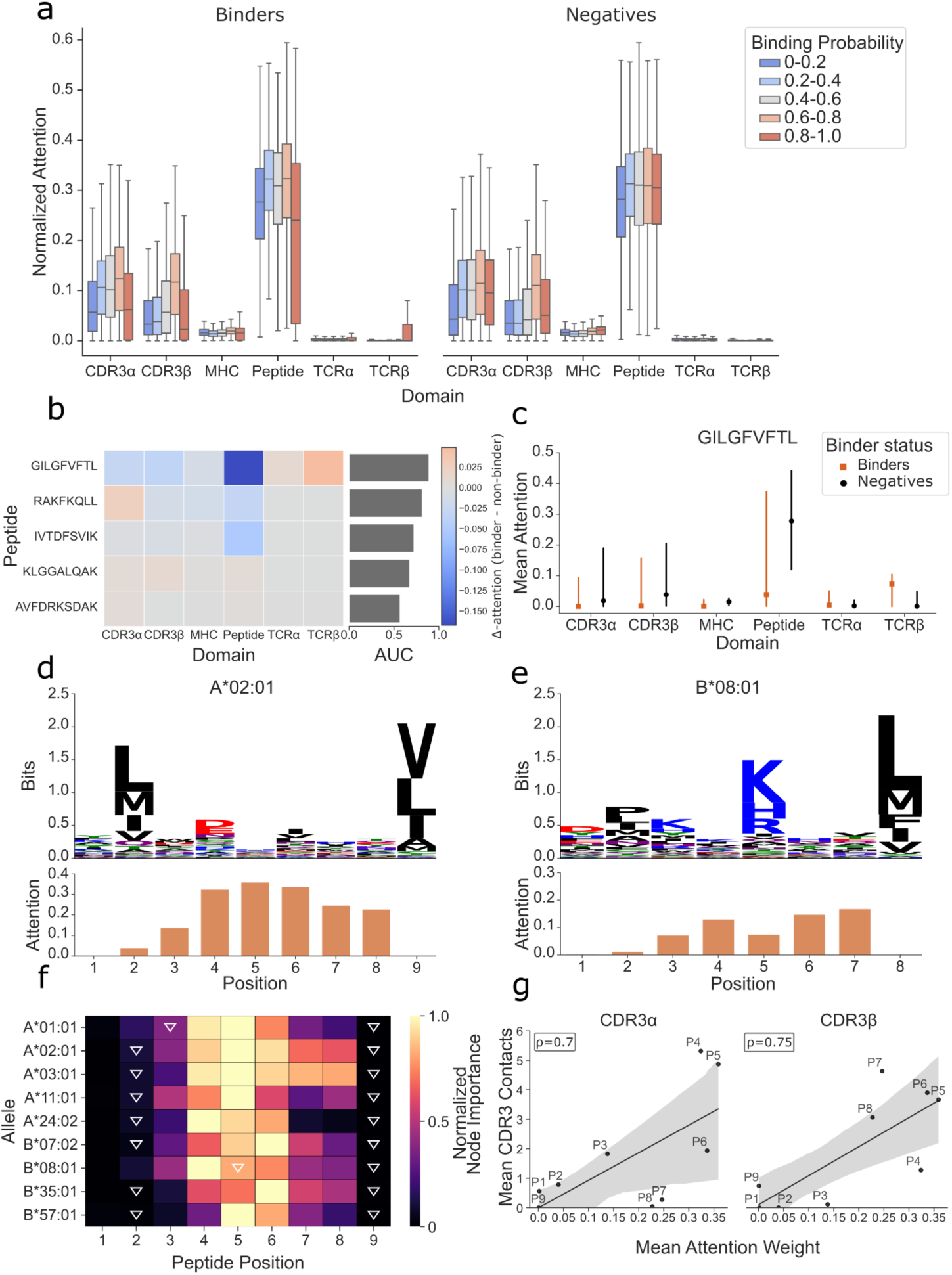
Domain and peptide-level attention patterns learned by t2pmhc-GCN reflect known determinants of TCR-pMHC recognition. (a) Domain-wise distribution of Min-Max normalized node attention weights derived from t2pmhc-GCNs attention-based global pooling, stratified by predicted binding probability quantiles for binders (left) and negatives (right). Attention is shown for the CDR3α, CDR3β, MHC, peptide, TCRα and TCRβ domains. (b) Heatmap of peptide-wise differences in mean domain attention between binders and negatives (Δ-attention), shown for the five most frequent peptides in the test dataset. Domains where less attention is provided to binders compared to negatives are depicted in blue, while domains where more attention is provided to binders appear in red. The accompanying barplot on the right side of the heatmap shows the AUC predictive performance achieved by t2pmhc-GCN for the respective peptide. (c) Domain-wise attention profiles for binders (orange) and negatives (black) for the peptide GILGFVFTL. (d,e) Comparison of position-specific peptide attention with MHC binding constraints. For A*02:01 9-mers (d) and B*08:01 8-mers (e), sequence logo based on the MHC Motif Atlas^58^ summarizing the allele-specific peptide-motifs (top), and mean attention assigned to each peptide position (bottom) are shown. (f) Heat map of mean position-wise peptide attention across MHC alleles with sufficient binder counts (>100 samples). Canonical MHC anchor positions derived from the MHC Motif Atlas are indicated by white triangles. (g) Correlation analysis in TCR-pMHC binder samples of mean attention weight assigned by t2pmhc-GCN to the individual peptide positions (P1-P9) in TCR-pMHC binders and the mean number of contacts formed with CDR3α (left) and CDR3β (right) in A*02:01-restricted peptides in the combined test set, with linear regression fits and 95% confidence intervals. Abbreviations: TCR, T cell receptor; MHC, major histocompatibility complex; CDR3, complementarity-determining region 3; GCN, graph convolutional network; GAT, graph attention network.

This effect was further explored for peptides in the test sets with sufficient sample size (> 1,000 observations) by computing the domain-wise attention differences between binders and negatives (Δ-attention) (Fig. 4b). This revealed a distinct pattern for the peptide GILGFVFTL, which achieved high predictive performance across both datasets (AUC=0.89). The observations for this peptide (Fig. 4c) correspond to the aforementioned pattern. Binders exhibited reduced peptide attention, accompanied by a slight increase in TCRβ attention and small decreases in CDRα and CDRβ attention. Hierarchical clustering of the Δ-attention identified a cluster of peptides displaying similar attention redistribution patterns with a focus on decreased peptide attention in binders (Supplementary Fig. 1d). Peptides in this cluster binding the A*02:01 allele showed consistently strong predictive performance.

Together, these findings indicate that t2pmhc-GCN not only assigns attention to biologically relevant regions but can also adapt its attention distribution in an allele- and peptide-specific manner.

### t2pmhc-GCN highlights biologically meaningful residues within the peptide sequence

t2pmhc-GCN attention focuses on the peptide domain, receiving 3.68-fold more attention than the next-highest domain (CDR3a). This strong focus on the peptide domain raises the question of how attention is distributed across individual peptide residues. To address this, we analyzed all binder samples containing 9-mer peptides from the immrep23 and the public test set. Mean attention values were computed per peptide position and compared to established MHC-binding motifs as described in the MHC Motif Atlas^58^ (Fig. 4d). Each MHC allele with a sufficiently large sample size (> 100 binders) was analyzed independently.

MHC anchor residues were defined as the two most conserved peptide positions in the MHC Motif Atlas. Conservation was determined based on the computed per-position information content in bits. All alleles in the datasets exhibited an anchor at position 9 (P9). t2pmhc-GCN consistently assigned very low attention to this position across alleles (mean = 0.00098 ± 0.0016), indicating that the model correctly identifies P9 as an MHC-binding anchor rather than a TCR-interacting residue. Most alleles also contained a second anchor at P2, which similarly received low attention (mean = 0.076 ± 0.05).

An exception is the allele B*08:01. This allele has an anchor at P5 which t2pmhc-GCN did not downweight (mean attention: 0.82). B*08:01 primarily binds 8-mer peptides, which is reflected in a large imbalance between 8-mer (6010 samples) and 9-mer (369 samples) peptides in the training data for this allele. Considering only 8-mers, downweighting of the anchor at P5 was observable (mean: 0.075 ± 0.13) (Fig. 4e).

For most alleles, the highest attention was assigned to positions P3-P8 (P3: 0.28 ± 0.14, P4: 0.79 ± 0.23 P5: 0.92 ± 0.11, P6: 0.84 ± 0.13, P7: 0.46 ± 0.23, P8: 0.39 ± 0.23), which are less constrained by MHC binding and thus more likely to participate in TCR recognition (Fig. 4f). To further validate this observation, we analyzed residue-level contact patterns focusing on the TCR-pMHC binders of the A*02:01 allele, accounting for allele-specific anchor positions. We found that P4 and P5 made the most contacts with CDR3α, while P5-P8 interacted most frequently with CDR3β in both binders and negative. In contrast, P1, P2 and P9 exhibited minimal contact with either CDR3 loop (Supplementary Fig. 2a). These findings are consistent with a recent STCRDab-based study^59^. Furthermore, we found a strong positive correlation between attention and CDR3 contacts (CDR3α: Spearman ρ = 0.70, CDR3β: Spearman ρ = 0.75) (Fig. 4g). Conversely, positions with higher MHC contact frequencies (P1, P2, P9) showed a strong negative correlation with attention (Spearman(ρ) = -0.92) (Supplementary Fig. 2c). A partially distinct pattern emerged for A*02:01 negatives (Supplementary Fig. 2b). While the correlation of CDR3β contacts to attention weight remained strong (Spearman(ρ)=0.75), the association for CDR3α was reduced (Spearman ρ = 0.55), largely due to decreased attention at P4. Similarly, the negative correlation between MHC contacts and peptide attention weights was a little weaker (Spearman ρ = -0.85) (Supplementary Fig. 2d,e,f).

Collectively, these findings demonstrate that t2pmhc-GCN assigns attention in a manner that is consistent with known structural and biological principles of TCR-pMHC recognition, highlighting its ability to capture biologically meaningful residue-level interaction patterns.

### Integration of complex-wide structural information enables improved prediction of unseen peptides

Across three independent benchmark datasets, both t2pmhc variants demonstrated consistently strong performance on seen peptides, achieving AUCs comparable to or exceeding established sequence-based predictors (Fig. 5a,c,d, Supplementary Fig 3). On the Public Test Set and the epytope-viral dataset, t2pmhc-GAT achieved the highest AUC, outperforming all competing models, while t2pmhc-GCN showed competitive performance. On the immrep23 dataset, MixTCRpred variants performed best, although both t2pmhc models remained close in performance. t2pmhc exhibited robust predictive accuracy across datasets despite variability in dataset composition.

**Figure 5:**
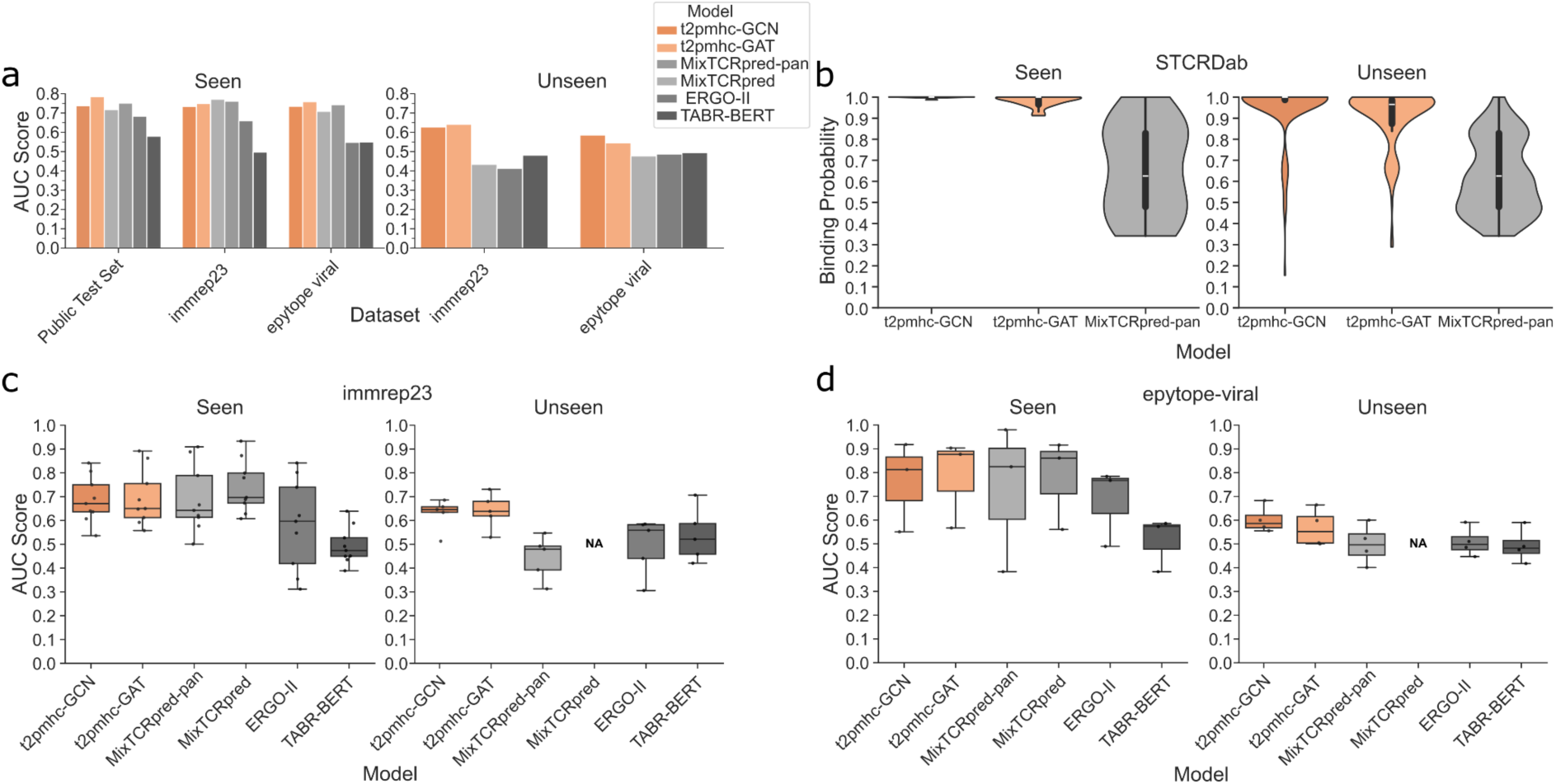
Performance comparison of t2pmhc and sequence-based predictors on seen and unseen peptide benchmarks. (a) Mean AUC scores across three benchmark datasets for seen (left) and unseen peptides (right). (b) Calibrated binding probability distributions on crystallographic TCR-pMHC complexes from STCRDab^53^ for seen (left) and unseen peptides (right), comparing t2pmhc-GCN, t2pmhc-GAT and MixTCRpred-pan. (c, d) Peptide-level AUC scores on the (c) immrep23 and (d) epytope-viral datasets for seen (left) and unseen peptides (right). Abbreviations: STCRDab: Structural T-Cell Receptor Database

Previous benchmarks have shown that prediction of unseen peptides remains challenging^3,52^. To evaluate the effect of structural information of the entire TCR-pMHC complex on improving generalization, we compared t2pmhc-GCN and t2pmhc-GAT against TABR-BERT, ERGO-II and MixTCRpred-pan on the immprep23 and the epytope-tcr viral datasets (Fig. 5a). MixTCRpred was not included in this benchmark as it cannot predict TCR-pMHC binding for unseen peptides. TABR-BERT, which performed worst on all seen datasets, ranked third on both unseen datasets. On the immrep23 dataset, t2pmhc-GAT showed the highest performance (AUC=0.642), followed by t2pmhc-GCN (AUC=0.626). TABR-BERT (AUC=0.481) showed near random performance, while MixTCRpred-pan (AUC=0.434) and ERGO-II (AUC=0.413) performed even poorer on this dataset. Notably, the t2pmhc model variants were the only models tested here to achieve peptide-level AUC values above 0.5 across all peptides in the immrep23 dataset (t2pmhc-GAT: mean AUC=0.639±0.08; t2pmhc-GCN: mean AUC=0.627±0.07) (Fig. 5c).

The epytope-viral dataset shows a similar pattern. Here, t2pmhc-GCN showed the strongest performance (AUC=0.586), followed by t2pmhc-GAT (AUC=0.545). TABR-BERT (AUC=0.495), ERGO-II (AUC=0.488) and MixTCRpred-pan (AUC=0.477) again demonstrated near-random performance on this dataset. The t2pmhc variants were the only models capable of achieving AUCs > 0.5 for all peptides in the dataset (t2pmhc-GCN: mean AUC=0.603±0.06, t2pmhc-GAT: mean AUC=0.567±0.08) (Fig. 5d).

Together, these results demonstrate that incorporating complex-wide structural information can improve generalization to unseen peptides.

### t2pmhc demonstrates accurate binding prediction with high quality structures

Structure prediction poses a substantial challenge for approaches such as t2pmhc, where performance is tightly linked to the success of the underlying structure prediction. To evaluate the predictive capabilities of the t2pmhc model variants under ideal structural conditions, we used crystallized TCR-pMHC structures from STCRDab as model inputs. Performance was assessed using the binding probabilities of t2pmhc-GCN, t2pmhc-GAT and MixTCRpred-pan. Binder prediction probability distributions were placed on a common scale using isotonic calibration, enabling direct comparison of probabilities between models (Supplementary Fig. 4a,b). For seen peptides, both t2pmhc variants produced binding probabilities strongly concentrated near 1.0. t2pmhc-GCN predicted a binding probability > 0.9 in 100% of samples, while t2pmhc-GAT predicted a binding probability > 0.9 in 96% of samples.

For unseen peptides, the t2pmhc models exhibited similarly high binder confidence. t2pmhc-GCN predicted a binding probability > 0.9 in 81.6% of samples, and only 5.3% of samples received probabilities < 0.5. t2pmhc-GAT assigned a binding probability > 0.9 in 71.1% of samples, also assigning probabilities < 0.5 in only 5.3% of samples.

In comparison, MixTCRpred-pan produced substantially lower binding probabilities for both seen and unseen peptides. No samples received a binding probability > 0.9 and 60% (seen) and 73.7% (unseen) of samples were assigned probabilities < 0.5 (Fig. 5b, Table 2).

**Table 2:**
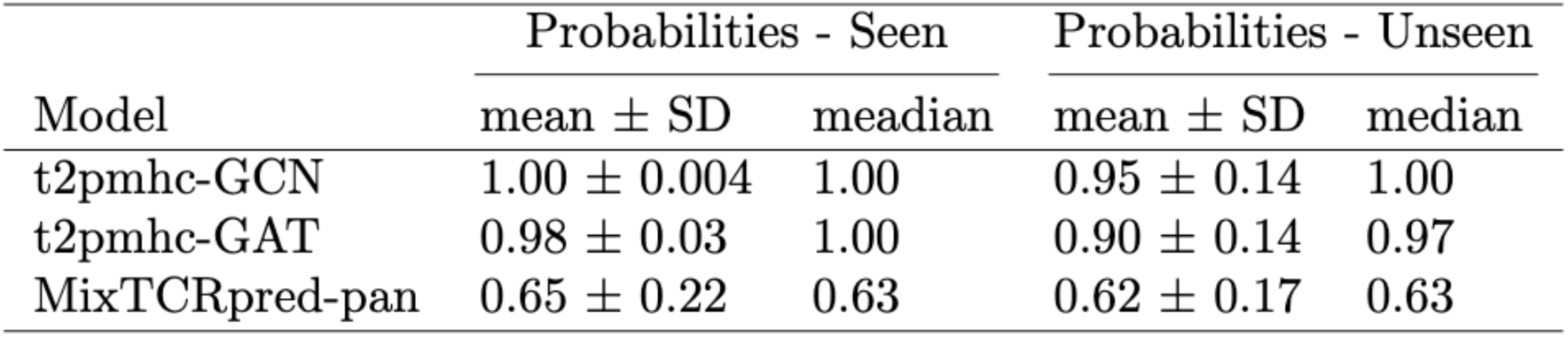
Binding probabilities of t2pmhc-GCN, t2pmhc-GAT and MixTCRpred-pan on the STCRDab dataset.

These results indicate that t2pmhc correctly identifies the vast majority of binders when provided with accurate structural input, even for unseen peptides, and despite being trained under suboptimal conditions on predicted structures, highlighting the potential of structure-based models and reinforcing the importance of accurate data inputs.

## Discussion

Many T cell-based immunotherapeutic strategies, comprising vaccination approaches, are usually developed without explicitly considering the binding of TCRs to peptide antigens presented by MHC molecules. While numerous TCR-pMHC binding predictors have been proposed in recent years, most rely primarily on sequence-level information^12–23^, with only a few incorporating structural information of the interaction^24–28^.

Contact map-based graph neural networks have recently been applied to structure-based TCR-pMHC binding prediction^24–27^, as well as to related tasks such as drug-target affinity prediction or protein-protein interactions^60,61^. For instance, STAG^24^ represents residues at the TCR-pMHC interaction site as a graph.

In this work we introduced t2pmhc, a structure-based deep learning framework that models residue-level graphs of the entire TCR-pMHC complex and systematically evaluated both GCN and GAT architectures on identical structural inputs.

In accordance with findings demonstrating the importance of the entire TCR-pMHC complex for binding^2^, TCR-pMHC complex structures were predicted using TCRdock^33^, a protein structure predictor specialized for this task. TCRdock constrains TCR docking orientation to native-like geometries by replacing AlphaFold Multimers MSAs with templates from ground truth TCR-pMHC structures^33^. Analysis of learned attention patterns revealed that t2pmhc-GCN captures biologically meaningful determinants of TCR-pMHC binding. At the domain level, attention was predominantly assigned to the peptide and CDR3 regions, consistent with their central role in antigen recognition. At the peptide residue level, canonical MHC anchor positions (P1, P2 and P9), which are crucial for peptide-MHC binding stability^62–64^ rather than TCR recognition, were consistently downweighted, while peptide positions linked to TCR binding (P5-P8)^59^ received high levels of attention. These findings indicate that the model implicitly differentiates residues involved in MHC binding from those mediating TCR contact. Given that attention mechanisms do not necessarily provide model explanations^65,66^, the strong agreement between attention patterns and independent structural evidence, particularly the correlation with CDR3 contact frequencies and the downweighting of canonical anchor residues, suggests that t2pmhc-GCN captures biologically meaningful features that drive its predictions and provides insight into model focus.

Beyond global trends, t2pmhc-GCN learned domain-, allele- and peptide-dependent attention patterns, reflecting the known variability in TCR-pMHC binding modes^4^. Attention redistributions were observed in high-confidence binders for specific alleles and even for individual peptides. For the A*02:01 allele, a cluster of peptides exhibited similar attention redistributions patterns, suggesting a shared allele-specific recognition mode. In these cases, increased attention was observed in domains that are not typically dominant or linked to TCR-pMHC recognition. For example, TCR binders of the GILGFVFTL peptide showed an increase in attention in the TCRβ domain. These findings further emphasize the importance of including the entire TCR-pMHC complex, as such non-canonical, context-specific recognition patterns would otherwise be inaccessible to methods focusing on isolated chains or domains. A comparable level of biologically consistent attention patterns was not observed for t2pmhc-GAT. Here, attention was preferentially assigned to large structural domains, namely the MHC, TCRα and TCRβ chains, potentially reflecting graph topology biases or degree effects.

Nevertheless, t2pmhc-GAT demonstrated comparable and often superior predictive performance to t2pmhc-GCN. This may reflect the increased expressiveness of attention-based message passing, which integrates edge features directly into the multi-head attention mechanism. In addition, the adaptive, per-edge weighting of interactions may enable the model to better accommodate structural and compositional heterogeneity.

These observations highlight a practical trade-off between predictive performance, biological consistency, and computational efficiency. While t2pmhc-GAT yields stronger classification results and thus is preferable in settings where classification is the primary objective, t2pmhc-GCN offers faster inference and yields attention distributions that align more closely with known structural features of TCR-pMHC recognition at both domain and residue resolution. This underscores how architectural choices, even when applied to identical structural inputs, can substantially influence model behaviour.

Both t2pmhc variants showed improved generalization to unseen peptides compared to state-of-the-art sequence-based methods^13–15^. The strong performance of MixTCRpred^15^ on the immrep23 dataset may reflect differences in dataset composition and is consistent with a recent benchmark^52^. The presented results highlight the importance of assessing benchmarks across multiple independent datasets, given the great fluctuation in performance of TABR-BERT, where the AUC on the best dataset was 16.6% higher than that on the worst dataset.

Notably, only the t2pmhc variants consistently achieved an AUC value > 0.5 across all evaluated test datasets. This suggests that the structural context of the full TCR-pMHC complex provides information that cannot be inferred from sequence alone, reinforcing the notion that TCR-pMHC binding is fundamentally a structural problem. It should be noted, however, that datasets containing significant numbers of unseen peptides are rare and more such datasets will be necessary to confirm this trend. The demonstrated ability of t2pmhc to generalize to unseen peptide samples of high structural quality is particularly relevant in the context of peptide vaccination^39,67,68^, or similarly mRNA vaccinations, where candidates are, by definition, novel and absent from training data. In this setting, the high consistency of the attention mechanisms of t2pmhc-GCN with biological concepts of TCR-pMHC binding can provide a foundation for integrating TCR sequencing data into peptide prioritization workflows, offering both predictive power and mechanistic insight into model decisions. Likewise, we anticipate that t2pmhc will enable facilitated immunomonitoring using TCR sequencing.

Evaluation of the t2pmhc variants on accurate crystallographic structures from STCRDab provided insights into their upper bound performance. When provided with accurate structures, t2pmhc produced near-deterministic binding predictions for both seen and unseen peptides, whereas the sequence-based MixTCRpred-pan yielded performance comparable to that of the other test sets. These findings suggest that current performance limitations of approaches like t2pmhc are largely driven by the quality of structure prediction rather than by model quality. This experiment was necessarily restricted to positive samples and relied on calibrated predicted binding probabilities rather than AUC values. Nonetheless, it indicates that improvements in structure prediction are likely to translate into improved binding prediction performances by the t2pmhc variants. Recent advancements in this field, particularly the emergence of AlphaFold 3^31,69^ and new specialized TCR-pMHC binding predictors and benchmarks^70,71^, raise optimism that higher-quality TCR-pMHC complex structures will soon become available, enabling more comprehensive and quantitative evaluation, including the incorporation of reliable negative samples.

Next to structural uncertainty, a broader limitation of the field remains the scarcity of large-scale, high-quality TCR-pMHC binder data. Since new models are likely to rapidly incorporate new data into their training, the number of unseen peptides remains low, complicating the evaluation of models for this problem.

As more data become available, both t2pmhc variants are expected to become more robust, and the better performing t2pmhc-GAT architecture may also improve in terms of biologically consistent attention patterns.

t2pmhc is a structure-based framework for TCR-pMHC binding prediction matching state-of-the-art sequence-based methods on seen peptides while outperforming them on the more challenging task of predicting binding to unseen peptides. t2pmhc is distributed via an easy-to-use command-line interface and Docker container to ensure usability and reproducibility. Analysis of learned attention patterns shows that t2pmhc-GCN prioritizes biologically relevant regions while downweighting canonical MHC anchor residues. Our results demonstrate that, when provided with accurate structures, t2pmhc can predict binders with very high confidence, highlighting its potential to be used to integrate TCR sequencing information into peptide antigen prioritization pipelines for T cell-based immunotherapy approaches. Ultimately, this approach may contribute to more informed immunotherapeutic design and improved patient care.

## Data availability

TCR-pMHC samples were retrieved from the VDJdb (https://vdjdb.com/), the McPAS-DB (https://friedmanlab.weizmann.ac.il/McPAS-TCR/) and the IDEB (https://www.iedb.org/).

## Code availability

t2pmhc is available on GitHub (https://github.com/qbic-pipelines/t2pmhc). The repository contains a Docker container to run the model reproducibly as well as the training data. The benchmarking pipeline is available on GitHub (https://github.com/qbic-pipelines/t2pmhc_benchmark). The pipeline to predict the structures is available on GitHub (https://github.com/mapo9/nf-core_proteinfold/tree/tcrdock).

## Acknowledgements

We thank M. Seybold from the Quantitative Biology Center and B. Gläßle from the M3 Research Center for their excellent technical support.

## Author Contributions

M.P., S.N. and J.S.W. conceptualized this study. S.N., J.S.W, M.D., A.N. and J.B. supervised this study. M.P. curated the data, developed the models, performed the bioinformatics analyses and benchmark. E.B. integrated TCRdock into nf-core/proteinfold. M.P, S.N. and J.S.W. wrote the manuscript. A.N., J.B., J.Sc., M.D., J.St. provided feedback on the manuscript. M.C., M.D., J.Sc. and J.St. gave continuous feedback during the study. S.N. and J.S.W. provided funding for the study.

## Author Information

Sven Nahnsen and Juliane S. Walz contributed equally to this work.

### Authors and Affiliations

**Quantitative Biology Center (QBiC), University of Tübingen, Tübingen, Germany**

Mark Polster, Josua Stadelmaier, Jonas Scheid, Marissa Dubbelaar & Sven Nahnsen

**Department of Peptide-based Immunotherapy, Institute of Immunology, University and University Hospital Tübingen, Tübingen, Germany**

Mark Polster, Jonas Scheid, Marissa Dubbelaar, Annika Nelde, Jens Bauer & Juliane S. Walz

**Cluster of Excellence iFIT (EXC2180) "Image-Guided and Functionally Instructed Tumor Therapies", University of Tübingen, Tübingen, Germany**

Jonas Scheid, Marissa Dubbelaar, Annika Nelde, Jens Bauer & Juliane S. Walz

**Clinical Collaboration Unit Translational Immunology, Department of Internal Medicine, University Hospital Tübingen, Tübingen, Germany**

Juliane S. Walz

**German Cancer Consortium (DKTK) and German Cancer Research Center (DKFZ), partner site Tübingen, Germany**

Juliane S. Walz, Jens Bauer

**Institute for Bioinformatics and Medical Informatics (IBMI), University of Tübingen, Tübingen, Germany**

Mark Polster, Elias Ball, Josua Stadelmaier, Jonas Scheid, Manfred Claassen & Sven Nahnsen

**Department of Computer Science, University of Tübingen, Tübingen, Germany**

Mark Polster, Elias Ball, Josua Stadelmaier, Jonas Scheid, Manfred Claassen & Sven Nahnsen

