## Supplementary Information for "t2pmhc: A Structure-Informed Graph Neural Network to Predict TCR-pMHC Binding"

for

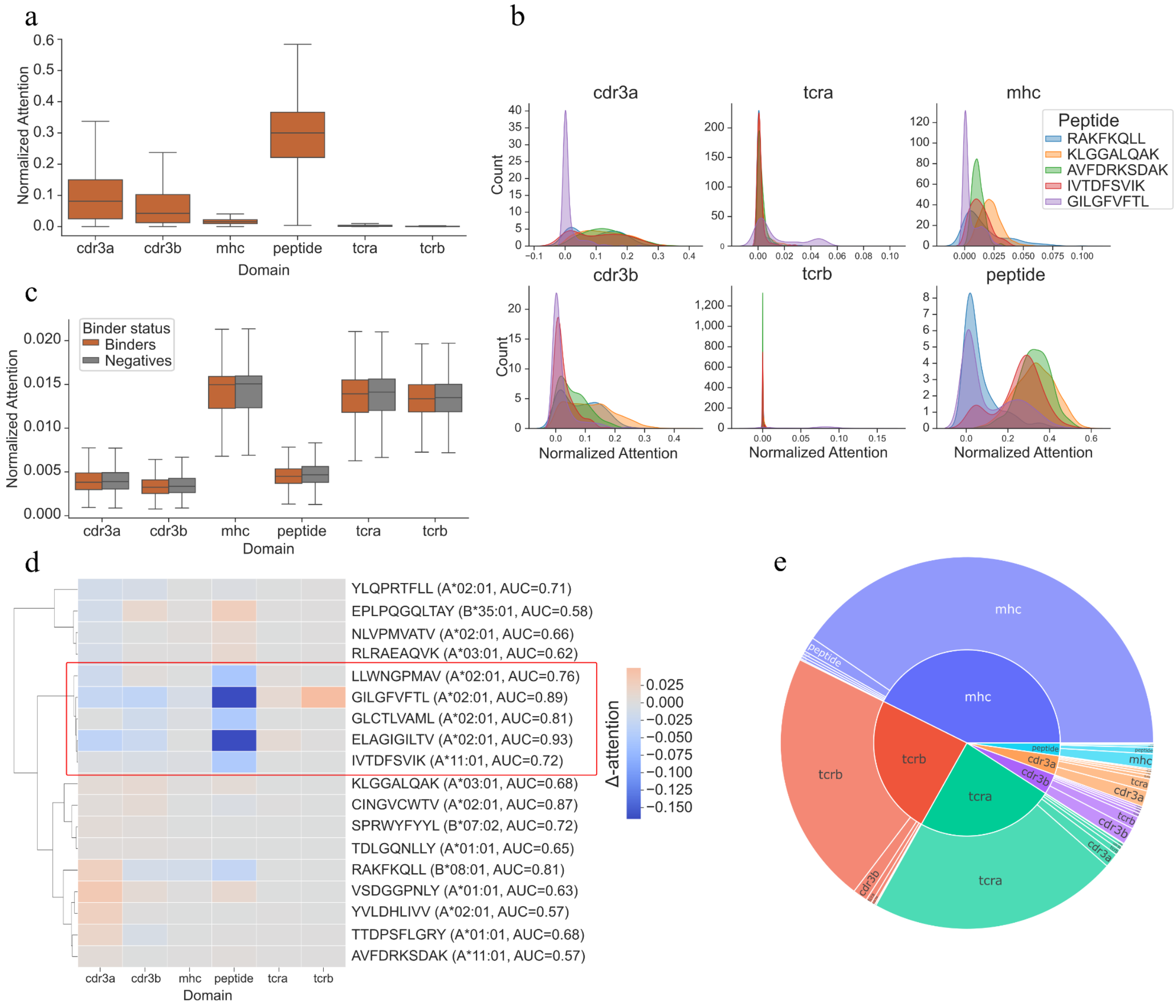


**Supplementary Fig. 1: Domain and peptide-specific attention patterns.**

(a) Domain-wise distribution of normalized node attention weights derived from t2pmhc-GCN, obtained from the attention-based global pooling layer. (b) Kernel density estimates of domain-wise t2pmhc-GCN attention weight distribution for the five most abundant peptides in the combined test set (immrep23 and public test set), shown as one density per peptide within each domain. (c) Domain-wise distribution of normalized node attention weights derived from t2pmhc-GAT, stratified by binder status (Binders (orange) and Negatives (grey)). (d) Hierarchical clustering of peptide-wise differences in mean domain attention between binders and negatives (Δ-attention) for peptides with sufficient sample size (> 1,000 samples) in the t2pmhc-GCN model. Blue indicates domains receiving less attention in binders than in negatives, while red indicates domains receiving more attention in binders than in negatives. The AUC achieved using t2pmhc-GCN for this peptide in the combined test set, as well as the MHC allele to which the respective peptide binds, is shown. (e) Sunburst plot summarizing the attention flow between domains in t2pmhc-GAT. The inner ring denotes the source and the outer ring the receiving domain. Segment size corresponds to the magnitude of attention flow between the respective domains. Values are aggregated across samples in the combined test set.

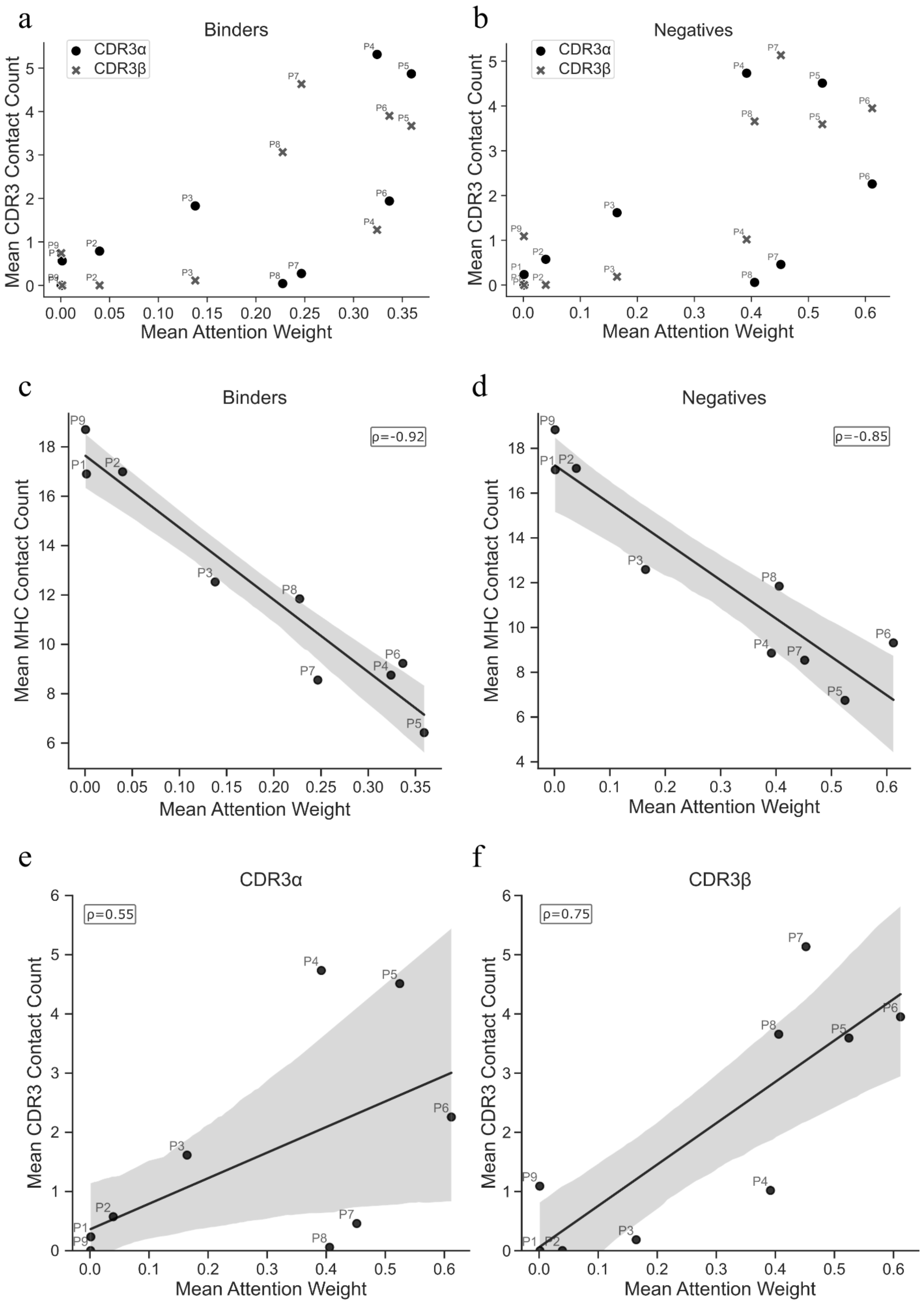


**Supplementary Fig. 2: Relationship between peptide attention and structural contact patterns in A*02:01-restricted complexes.**

(a) Mean attention weight assigned by t2pmhc-GCN to the individual peptide positions (P1-P9) in TCR-pMHC binders versus the mean number of contacts formed with CDR3α (x) and CDR3β (o) across A*02:01-restricted peptides in the combined test set. (b) Mean attention weight assigned by t2pmhc-GCN to the individual peptide positions (P1-P9) in TCR-pMHC negatives versus the mean number of contacts formed with CDR3α (x) and CDR3β (o) across A*02:01-restricted peptides in the combined test set. (c) Correlation analysis of mean attention weight assigned by t2pmhc-GCN to the individual peptide positions (P1-P9) and the mean number of contacts formed with the MHC domain in TCR-pMHC binders within the A*02:01-restricted peptides in the combined test set, with linear regression fits and 95% intervals. (d) Correlation analysis of mean attention weight assigned by t2pmhc-GCN to the individual peptide positions (P1-P9) and the mean number of contacts formed with the MHC domain in TCR-pMHC negatives within the A*02:01-restricted peptides in the combined test set, with linear regression fits and 95% intervals. (e) Correlation analysis in TCR-pMHC negative samples of mean attention weight assigned by t2pmhc-GCN to the individual peptide positions (P1-P9) in TCR-pMHC negatives and the mean number of contacts formed with CDR3α in A*02:01-restricted peptides in the combined test set, with linear regression fits and 95% confidence intervals.

(f) Correlation analysis in TCR-pMHC binder samples of mean attention weight assigned by t2pmhc-GCN to the individual peptide positions (P1-P9) in TCR-pMHC negatives and the mean number of contacts formed with CDR3β in A*02:01-restricted peptides in the combined test set, with linear regression fits and 95% confidence intervals.

Abbreviations: MHC, major histocompatibility complex; CDR3, complementarity-determining region 3.

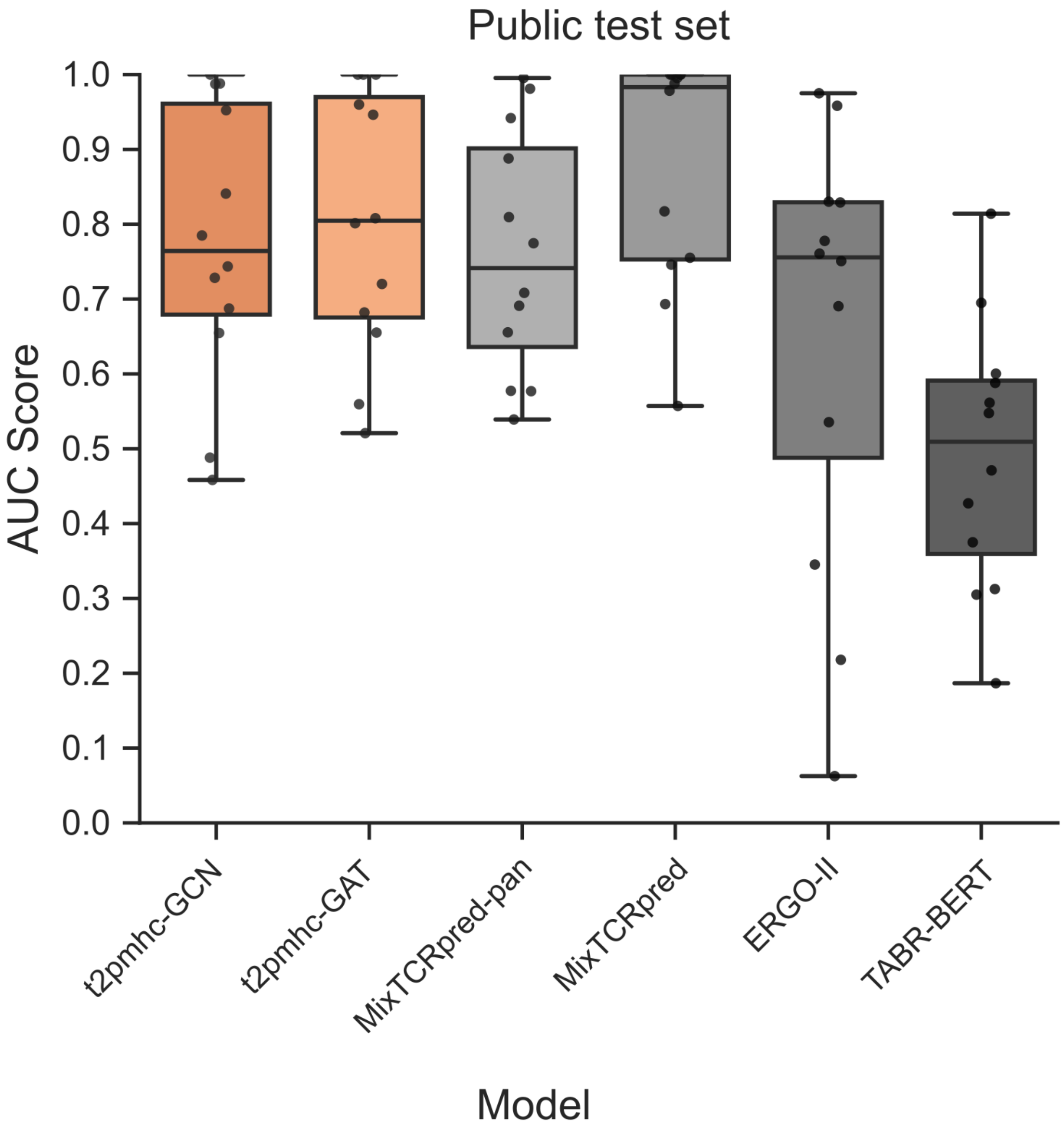


**Supplementary Fig. 3: Peptide-level AUC scores on the public test datasets for seen peptides. No unseen peptides were available for this dataset.**

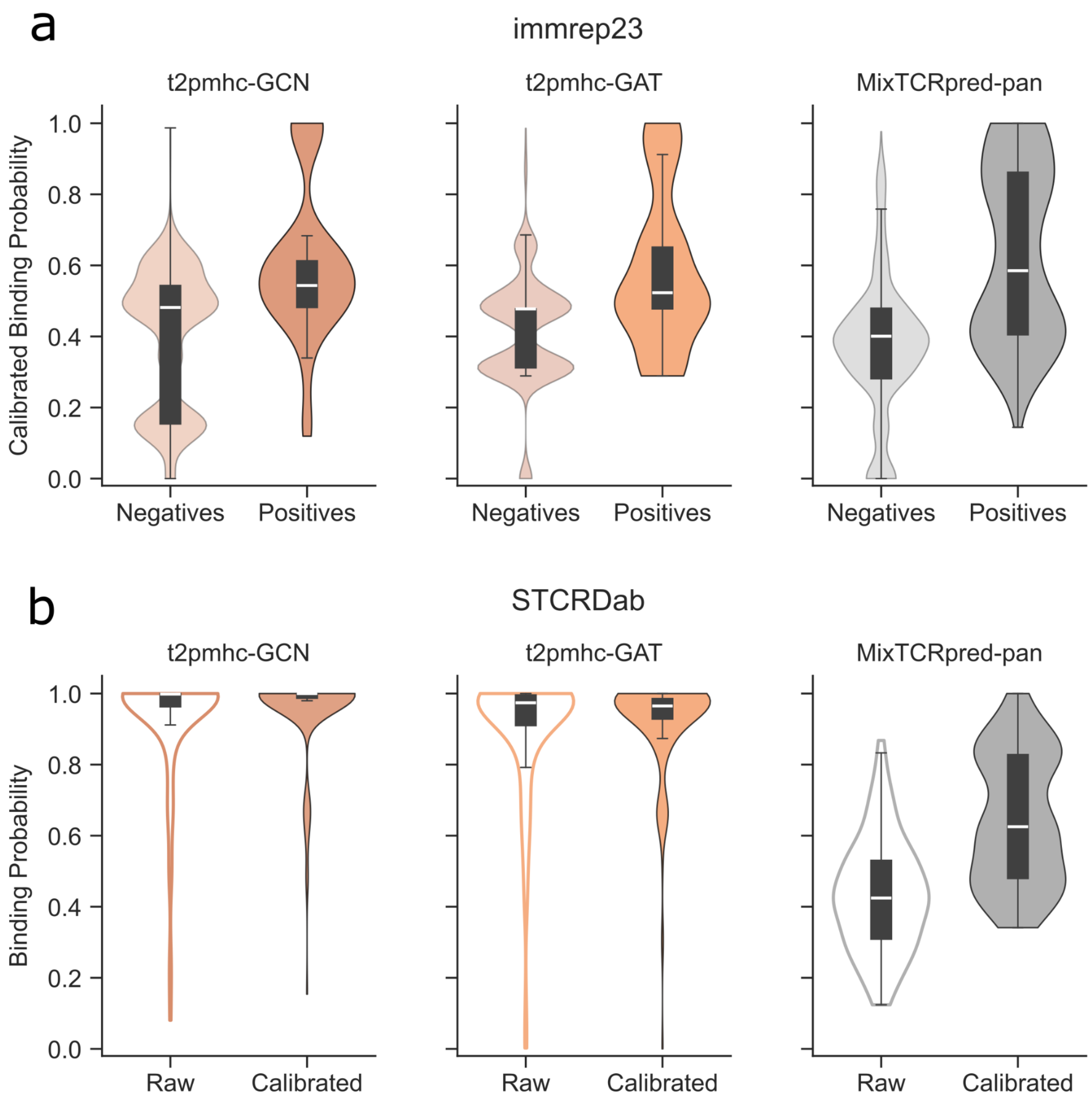


**Supplementary Fig. 4: Isotonic calibration validation across binding prediction models.** (a) Distribution of calibrated binding probabilities of negatives (light) and positives (dark) in the immrep23 test set, which served as the calibration training set for t2pmhc-GCN (left), t2pmhc-GAT (middle), and MixTCRpred-pan (right). Isotonic regression was fitted independently for each model to map raw predicted probabilities to calibrated values. (b) Distribution of raw (outline) and calibrated (filled) binding probabilities for TCR-pMHC binders from the Structural T-Cell Receptor Database (STCRdab) dataset for t2pmhc-GCN (left), t2pmhc-GAT (middle), and MixTCRpred-pan (right). The calibration was fitted based on the immrep23 test set as described in (a).

Abbreviations: STCRDab, The Structural T cell Receptor Database
